# CD40 and CD80/86 signaling in cDC1s mediate effective neoantigen vaccination and generation of antigen-specific CX3CR1^+^ CD8^+^ T cells in mice

**DOI:** 10.1101/2020.06.15.151787

**Authors:** Takayoshi Yamauchi, Toshifumi Hoki, Takaaki Oba, Kristopher Attwood, Xuefang Cao, Fumito Ito

## Abstract

The use of tumor mutation-derived neoantigen represents a promising approach for cancer vaccines. Preclinical and early-phase human clinical studies have shown the successful induction of tumor neoepitope-directed responses; however, overall clinical efficacy of neoantigen vaccines has been limited. One major obstacle of this strategy is the prevailing lack of sufficient understanding of the mechanism underlying the generation of neoantigen-specific CD8^+^ T cells. Here, we report a correlation between antitumor efficacy of neoantigen/toll-like receptor 3 (TLR3)/CD40 vaccination and the generation of antigen-specific CD8^+^ T cells expressing CX3C chemokine receptor 1 (CX3CR1) in a preclinical model. Mechanistic studies using mixed bone marrow chimeras identified that CD40 and CD80/86, but not CD70 signaling in Batf3-dependent conventional type 1 dendritic cells (cDC1s) is required for antitumor efficacy of neoantigen vaccine and generation of neoantigen-specific CX3CR1^+^ CD8^+^ T cells. Although CX3CR1^+^ CD8^+^ T cells exhibited robust *in vitro* effector function, depletion of this subset did not alter the antitumor efficacy of neoantigen/TLR3/CD40 agonists vaccination, suggesting that the expanded CX3CR1^+^ CD8^+^ T cell subset represents the post-differentiated *in vivo* effective CX3CR1-negative CD8^+^ T cell subset. Taken together, our results reveal a critical role of CD40 and CD80/86 signaling in cDC1s in antitumor efficacy of neoantigen-based therapeutic vaccines, and implicate the potential utility of CX3CR1 as a circulating predictive T-cell biomarker in vaccine therapy.

## Introduction

Despite objective therapeutic benefit seen in several clinical trials, the overall clinical success of the peptide-based vaccines has thus far been limited (1, 2). This is due, at least in part, to the low immunogenicity of self-antigens, the incomplete understanding of immunological mechanisms underlying effective priming of tumor-specific T cells, and the lack of a reliable biomarker for clinical response. Somatically mutated genes within tumors can generate neoantigens, which create *de novo* epitopes for T cells (3). Since neoantigens have not undergone central thymic selection, they can be deemed as highly immunogenic tumor-specific antigens (3). Recent advances of neoantigen identification by massive parallel sequencing and computational prediction of neo-epitopes have demonstrated that individualized mutanome-based vaccinations are promising (4–7).

The chemokine receptor, CX3CR1, was recently identified as a marker of effector T-cell differentiation (8, 9). Unlike other proliferation, co-stimulatory and co-inhibitory molecules such as Ki67, ICOS, PD-1 and CTLA-4 that are only transiently upregulated on T cells after activation, CX3CR1 is stably expressed on virus- and tumor-specific CD8^+^ T cells upon differentiation via unidirectional differentiation from the CX3CR1-negative (CX3CR1^−^) subset during the effector phase (8–10). Furthermore, our study and others have shown that CD8^+^ T cells expressing high levels of CX3CR1 exhibit decreased expression of L-selectin (CD62L) and CXCR3 (8–10), trafficking receptors necessary for entry across lymphoid organ high endothelial venules (HEV) and the tumor microvasculature, respectively (11, 12), and become more prevalent in peripheral blood (PB) at the end of the primary response (9, 10). These observations prompted us to hypothesize that effective vaccination would be associated with the increased frequency of antigen-specific CX3CR1^+^ CD8^+^ T cells in the periphery.

Here, we used a syngeneic mouse model of colon adenocarcinoma with the neo-epitope presented in major histocompatibility complex (MHC) class I H-2D^b^ molecules (13), and investigated the relationship between antitumor efficacy of neoantigen vaccination and the frequency of neoantigen-specific CX3CR1^+^ CD8^+^ T cells. Our results indicate that generation of circulating neoantigen-specific CX3CR1^+^ CD8^+^ T cells correlates with successful vaccination with mutated peptide and toll-like receptor 3 (TLR3)/CD40 agonists. Mechanistic studies using mixed bone marrow chimeras identified a key role of CD40 and CD80/86, but not CD70 signaling in Batf3-dependent conventional type 1 dendritic cells (cDC1s) for the generation of neoantigen-specific cytotoxic CX3CR1^+^ CD8^+^ T cells and therapeutic efficacy of neoantigen vaccine.

## Materials and Methods

### Mice

Female C57BL/6 mice were purchased from the Jackson Laboratory. CD40^−/−^ mice (B6.129S2-*Cd40lg^tm1Imx^*/J), CD80/86^−/−^ mice (B6.129S4-*Cd80^tm1Shr^ Cd86^tm2Shr^*/J), and Batf3^−/−^ mice (B6.129S(C)-*Batf3^tm1Kmm^*/J), CD2-Cre mice (C57BL/6-Tg(CD2-cre)1Lov/J), and CX3CR1-DTR mice (B6N.129P2-Cx3cr1^*tm3(DTR)Litt*^/J) on C57BL/6 background were purchased from the Jackson Laboratory, and bred in-house (Roswell Park Comprehensive Cancer Center). CD70^−/−^ mice on C57BL/6 background have been previously described (14). For inducible *Cd2-cre/Cx3cr1*^+/*DTR*^ mice, we crossed CD2-Cre mice with CX3CR1-DTR mice to generate *Cd2-cre/Cx3cr1*^+/*DTR*^ mice, allowing induction of DTR in CX3CR1^+^ CD8^+^ T cells. For depletion of CX3CR1^+^ CD8^+^ T cells, diphtheria toxin (Sigma) (250 ng/dose/mouse) was administered intraperitoneally (i.p.) every day starting from 1 day before neoantigen vaccination. All mice were 7 to 12 weeks old at the beginning of each experiment, and were maintained under specific pathogen-free conditions at the Roswell Park animal facility according to approved institutional guidelines.

### Cells

The murine B16F10 melanoma cell line was purchased from ATCC. The MC38 murine colon adenocarcinoma cell line was gift from Dr. Weiping Zou (University of Michigan). B16F10 and MC38 cells were maintained in RPMI (Gibco) supplemented with 10% FBS (Sigma), 1% NEAA (Gibco), 2 mM GlutaMAX-1 (Gibco), 100 U/ml penicillin-streptomycin (Gibco), and 55 μM 2-mercaptoethanol (Gibco). Cells were authenticated by morphology, phenotype and growth, and routinely screened for *Mycoplasma*, and were maintained at 37°C in a humidified 5% CO_2_ atmosphere.

### *In vivo* vaccination and cancer immunotherapy studies

Female C57BL/6 mice were injected s.c. with 5-8 × 10^5^ MC38 cells into the right flank. Mice were treated s.c. with 100 μg soluble Adpgk^Mut^ (ASMTNMELM) or AH1 (SPSYVYHQF) peptide (GenScript) with 50 μg poly(I:C) (InvivoGen) and 50 μg agonistic CD40 Ab (BioXCell, clone FGK45) in 100 μl of PBS into the left flank twice 1 week apart. PB, spleen and tumors were harvested for immune monitoring 1 week after the second vaccination. Tumor volumes were calculated by determining the length of short (*l*) and long (*L*) diameters (volume = *l*^2^ × *L*/2). Experimental end points were reached when tumors exceeded 20 mm in diameter or when mice became moribund and showed signs of lateral recumbency, cachexia, lack of response to noxious stimuli, or observable weight loss.

### Generation of bone marrow chimeras

To generate bone marrow chimeras, C57BL/6 mice were irradiated with 500 cGy followed by a second dose of 550 cGy 3 hours apart. To obtain donor bone marrow, femurs and tibiae were harvested and the bone marrow was flushed out. For Batf3^−/−^/WT, Batf3^−/−^/CD40^−/−^, Batf3^−/−^/CD80/86^−/−^, and Batf3^−/−^/CD70^−/−^ mixed bone marrow chimeras, 5 × 10^6^ bone marrow cells of a 1:1 mixture were injected to irradiated C57BL/6 WT mice. After 8-12 weeks, recipients were used for the experiments.

### Flow cytometry and intracellular granzyme, TNF-α and IFN-γ assay

Fluorochrome-conjugated antibodies are shown in Table S1. DAPI, LIVE/DEAD Fixable Aqua (Thermo Fisher Scientific) staining cells were excluded from analysis. For tetramer staining, PE-labeled MHC class I (H-2D^b^) specific for mouse Adpgk^Mut^ 299-307 (ASMTNMELM) (13) was kindly provided by the NIH Tetramer Core Facility. Detection of intracellular cytokines, granzyme A (GZMA), IFN-γ and TNF-α was performed as described before (10). For intracellular staining of IFN-γ and TNF-α, splenocytes and tumor-infiltrating cells were cocultured with 1 μmol/L of Adpgk^Mut^ or AH1 peptide in the presence of Brefeldin A (BD Biosciences) for 5h *in vitro*. Samples were analyzed using an LSR II or an LSRFortessa (BD Biosciences) with FlowJo software (TreeStar).

### Statistical Analysis

Continuous measures were compared between two groups using two-tailed t-tests (unpaired and paired) and between multiple groups using one-way ANOVA, with Tukey adjusted post-hoc tests. Survival outcomes were compared using the log-rank test (Mantel-Cox method). Correlations were assessed using the Pearson and Spearman correlation coefficients. Analyses were performed in GraphPad Prism 8.02 (GraphPad Software) and SAS v9.4 (Cary, NC) at a significance level of 0.05.

## Results

### Vaccination with neoantigen and TLR3/CD40 agonists elicits potent antigen-specific antitumor efficacy

To examine the relationship between antitumor efficacy of neoantigen vaccination and the generation of neoantigen-specific CD8^+^ T cells, we sought to validate a preclinical model of neoantigen vaccination using MC38 colon adenocarcinoma cells that harbor a single-epitope mutation within the Adpgk protein (**Supplemental Figure 1**) (13). In this model, to maximize the therapeutic efficacy of neoantigen vaccination with mutant Adpgk peptide (Adpgk^Mut^), dual TLR/CD40 stimulation was used as vaccine adjuvants **(Figure 1A**), which has been known to synergistically activate DCs, augment CD8^+^ T cell expansion, and mediate potent antitumor immunity in multiple preclinical models (14–20). Vaccination of Adpgk^Mut^, but not irrelevant AH1 peptide with agonistic anti-CD40 antibody (Ab), and a TLR3 agonist, poly(I:C) (referred to as Adpgk^Mut^/TLR3/CD40 hereafter) markedly delayed the growth of established MC38 tumors and improved survival **(Figure 1B**). Splenocytes and tumor-infiltrating cells from MC38 tumor-bearing mice treated with Adpgk^Mut^**/**TLR3/CD40 vaccination contained substantially higher frequency of CD8^+^ T cells that could produce IFN-γ and TNF-α against Adpgk^Mut^ but not irrelevant AH1 antigen **(Figure 1C** and **Supplemental Figure 2**), suggesting the generation of antigen-specific CD8^+^ T cells. These findings validate a preclinical model of neoantigen vaccine-based therapy to investigate the role of neoantigen-specific CD8^+^ T cells.

**Figure 1.**
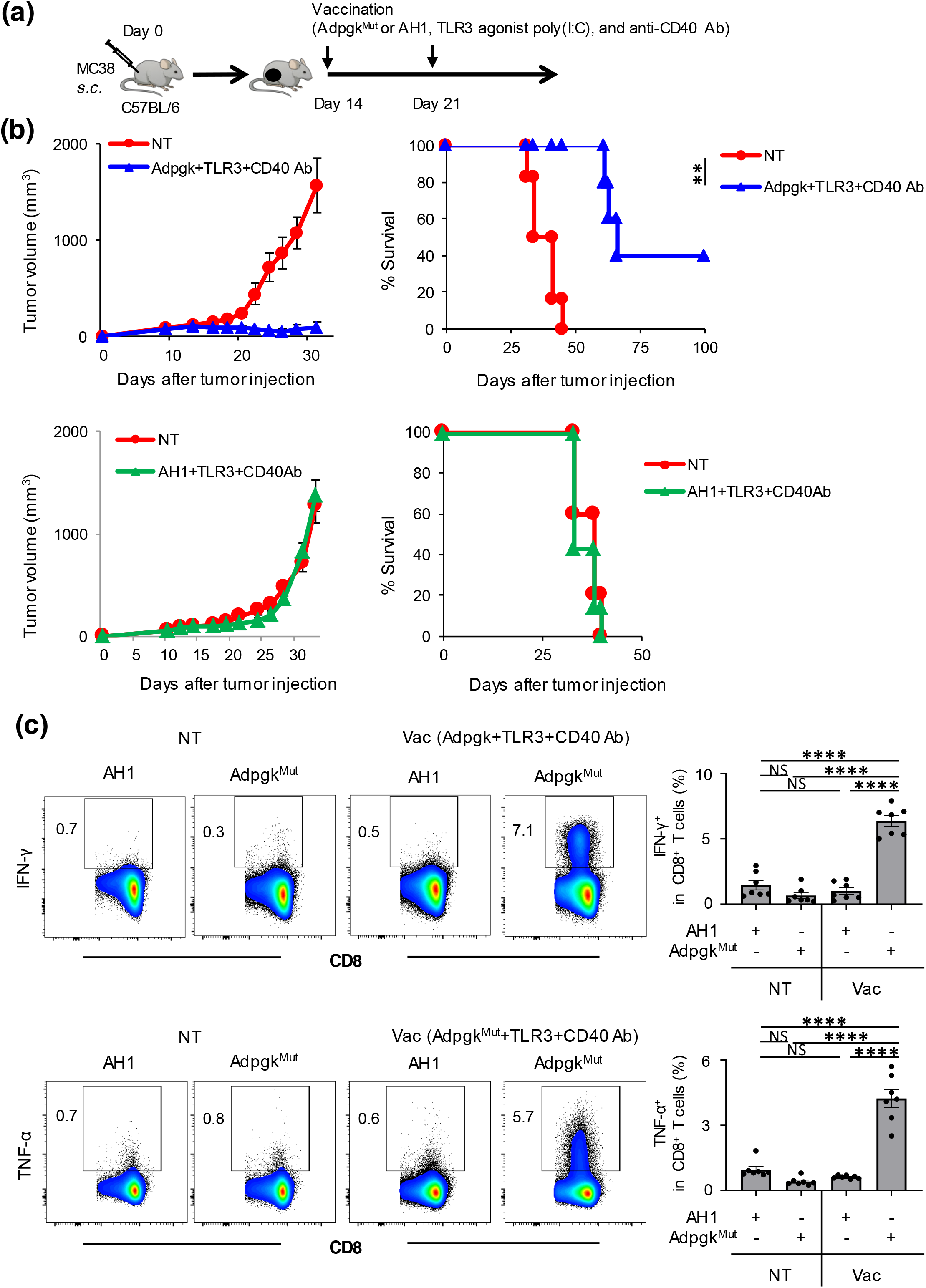
Vaccination with neoantigen and TLR3/CD40 agonists elicits potent antigen-specific antitumor efficacy. (**a**) Experimental scheme for evaluation of neoantigen/TLR3/CD40 agonists vaccination (Vac) in an MC38 tumor model (refer Materials and Methods for details). (**b**) Tumor growth and survival curves in MC38 tumor-bearing C57BL/6 mice treated with PBS (NT: non-treatment) or Adpgk^Mut^/TLR3/CD40 agonists vaccination (upper) (n = 6 mice per group) and PBS (NT) or AH1/TLR3/CD40 agonists vaccination (lower) (n = 5-7 mice per group). (**c**) Left shows representative FACS plots showing CD8 and IFN-γ (upper) or TNF-α (lower) expression gated with CD8^+^ T cells in splenocytes of MC38-tumor bearing mice 1 week after second PBS injection (NT) or Adpgk^Mut^/TLR3/CD40 agonists vaccination. Splenocytes were co-cultured with Adpgk^Mut^ or AH1 peptide in the presence of Brefeldin A for 5 hrs before intracellular staining. Numbers denote percentage of IFN-γ^+^ or TNF-α^+^ cells. Right panel shows the frequency of the IFN-γ^+^ or TNF-α^+^ cells in CD8^+^ T cells in each group (n = 7 mice per group). NS not significant, \*\*\*\**P* < 0.0001 by log-rank (Mantel-Cox) test (**b**) and one-way ANOVA with Tukey’s multiple comparisons (**c**). Mean ± SEM.

### Effective neoantigen/TLR3/CD40 stimulations correlate with generation of antigen-specific effector CX3CR1^+^ CD8^+^ T cells

We examined whether effective neoantigen vaccination could generate CD8^+^ T cells expressing CX3CR1, a marker of effector T-cell differentiation (8, 9). To this end, PB, spleen and tumors were harvested 1 week after second Adpgk^Mut^**/**TLR3/CD40 vaccination, and CX3CR1 expression was evaluated in CD8^+^ and neoantigen-specific CD8^+^ T cells using an Adpgk^Mut^-specific tetramer (Tet) (**Supplementary Figure 3**). Adpgk^Mut^**/**TLR3/CD40 vaccination substantially increased the frequency of CD8^+^ and Tet^+^ CD8^+^ T cells expressing CX3CR1 in PB and spleen while the frequency of the CX3CR1^+^ subset remained unchanged in tumors (**Figure 2A**).

**Figure 2.**
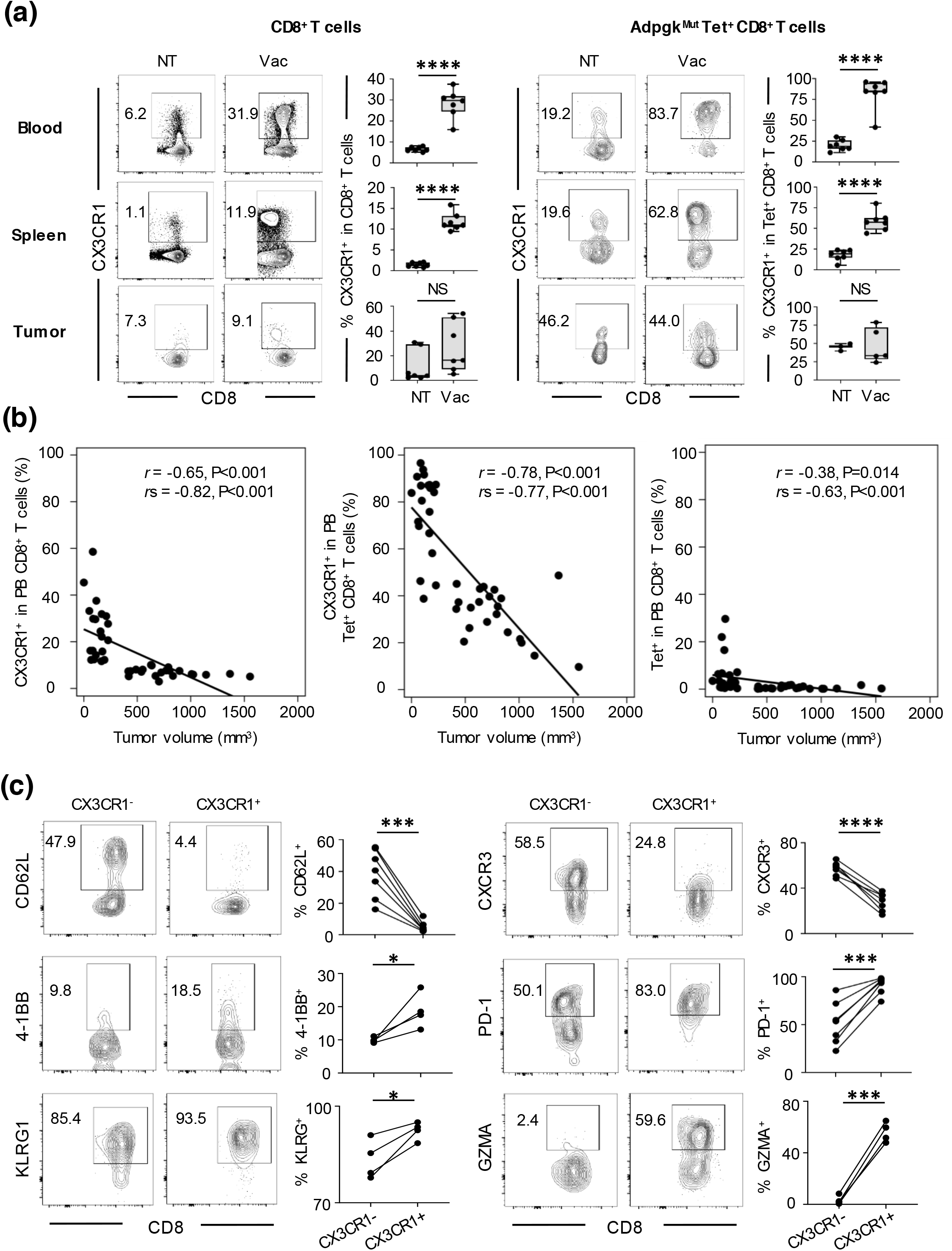
Effective neoantigen/TLR3/CD40 stimulations correlate with generation of antigen-specific effector CX3CR1^+^CD8^+^ T cells. (**a**) Representative FACS plots showing CX3CR1 expression in CD8^+^ (left) or Adpgk^Mut^ tetramer (Tet)^+^CD8^+^ T cells (right) in peripheral blood (PB) (upper), spleen (middle) and tumors (lower) from MC38-tumor bearing mice 1 week after second Adpgk^Mut^/TLR3/CD40 agonists vaccination (Vac). Numbers denote percentage of CX3CR1-positive cells. Frequency of CX3CR1-positive cells from the CD8^+^ or Tet^+^CD8^+^ T cells is shown in the right panels (n = 3-7 mice per group). Data shown are representative of three independent experiments. (**b**) Scatter plot of the frequency of PB CX3CR1^+^CD8^+^ (left), CX3CR1^+^Tet^+^CD8^+^ T cells (middle), and Tet^+^CD8^+^ T cells against tumor volume in untreated and treated mice 1 week after second Adpgk^Mut^/TLR3/CD40 agonists vaccination. Correlation is shown using Pearson correlation (*r*) and Spearman correlation coefficients (*r*_s_). Data shown are representative of two independent experiments. (**c**) Representative FACS plots of CX3CR1^−^ (left) or CX3CR1^+^ (right) cells gated with Tet^+^CD8^+^ T cells in PB of MC38-tumor bearing mice 1 week after second Adpgk^Mut^/TLR3/CD40 agonists vaccination. Numbers denote percentage of each marker-positive cells. Frequency of marker-positive cells from the CX3CR1^−^ or CX3CR1^+^ subset is shown in the right panels for each marker (n = 4-7 mice per group). Data shown are representative of two independent experiments. NS: not significant, \**P* < 0.05, \*\*\**P* < 0.001, \*\*\*\**P* < 0.0001 by unpaired (**a**) and paired two-tailed *t*-test (**c**). Mean ± SEM.

Previous clinical studies showed that functional assessment of T cells such as IFN-γ production and/or cytolytic activity was correlated with response in patients treated with peptide vaccines (21–23). Therefore, we evaluated CX3CR1 expression of T cells expressing effector cytokines, and found that the majority of CD8^+^ T cells producing effector cytokine against Adpgk^Mut^ were largely positive for CX3CR1 expression (**Supplementary Figure 4A**). Accordingly, frequency of the CX3CR1^+^ subset in circulating Tet^+^ CD8^+^ T cells correlated with the frequency of CD8^+^ T cells secreting IFN-γ^+^ and TNF-α^+^ against Adpgk^Mut^ antigen in spleen (**Supplementary Figure 4B**).

Next, we assessed relationship between tumor volume and antigen-specific CD8^+^ T-cell expansion and differentiation in PB from treated and untreated mice. There was a strong inverse correlation between frequency of circulating CD8^+^ and Tet^+^ CD8^+^ T cells expressing CX3CR1 and tumor volume while the frequency of total Tet^+^ CD8^+^ T cells without CX3CR1 criteria had a moderate inverse correlation with tumor volume **(Figure 2B**).

Phenotypical and functional evaluation of circulating Adpgk^Mut^-specific CX3CR1^−^ and CX3CR1^+^ subsets revealed that the CX3CR1^+^ subset contained significantly more 4-1BB^+^, PD-1^+^, KLRG^+^ and GZMA^+^ populations (**Figure 2C**), suggesting recently-activated effector T cells. However, CX3CR1^+^ CD8^+^ T cells exhibited decreased expression of trafficking molecules, CD62L and CXCR3, involved in T-cell access into lymphoid tissues and tumor microenvironment (TME), respectively (11, 12), in line with the higher frequency of the CX3CR1^+^ subset in PB in treated mice (**Figure 2A**).

### *In vivo* antitumor efficacy of antigen-specific CX3CR1+ CD8+ T cells elicited by neoantigen/TLR3/CD40 agonist vaccination

Previous preclinical studies using CX3CR1^−/−^ mice or CX3CR1 inhibitor suggested a significance of the CX3CR1/CX3CL1 axis in T cell/NK-cell mediated antitumor immunity (24–28). However, CX3CR1 is expressed not only on T and NK cells, but also on monocytes, DCs, monocytic myeloid-derived suppressor cells (M-MDSCs) and tumor-associated macrophages (TAMs) (29, 30), and antitumor efficacy of CX3CR1^+^ CD8^+^ T cells against established tumors remains elusive. Recently, we have reported negligible antitumor efficacy of CX3CR1^hi^ CD8^+^ T cells using a mouse model where tumor antigen-specific CX3CR1^hi^ CD8^+^ T cells can be selectively depleted *in vivo* with diphtheria toxin (DT) (10). Furthermore, adoptive transfer of CX3CR1^+^ CD8^+^ T cells did not affect tumor growth nor survival while the CX3CR1^−^ subset displayed potent antitumor efficacy (10).

To elucidate antitumor efficacy of CX3CR1^+^ CD8^+^ T cells elicited by neoantigen/TLR3/CD40 agonists vaccination, we crossed CD2-Cre mice with CX3CR1-DTR mice, and generated *Cd2-cre/Cx3cr1*^+/*DTR*^ mice. Expression of *Cd2-cre* excises the *loxP*-floxed stop cassette upstream of the DTR-coding region, allowing for DTR expression and CX3CR1^+^ CD8^+^ T cells can be depleted upon administration of DT, with no effect on CX3CR1^−^ CD8^+^ T cells (**Figure 3A, B**). MC38 tumor-bearing *Cd2-cre/Cx3cr1^+/DTR^* mice were treated with Adpgk^Mut^/TLR3/CD40 vaccination, and received phosphate buffer saline (PBS) or DT injections every day starting 1 day before vaccination (**Figure 3B**). We found that depletion of CX3CR1^+^ CD8^+^ T cells did not alter antitumor efficacy of neoantigen vaccination (**Figure 3C**). This is in line with our recent study using a mouse model of adoptive T-cell therapy (10), suggesting that the expanded CX3CR1^+^ CD8^+^ T cells represent the post-differentiated *in vivo* effective CX3CR1-negative CD8^+^ T cell subset.

**Figure 3.**
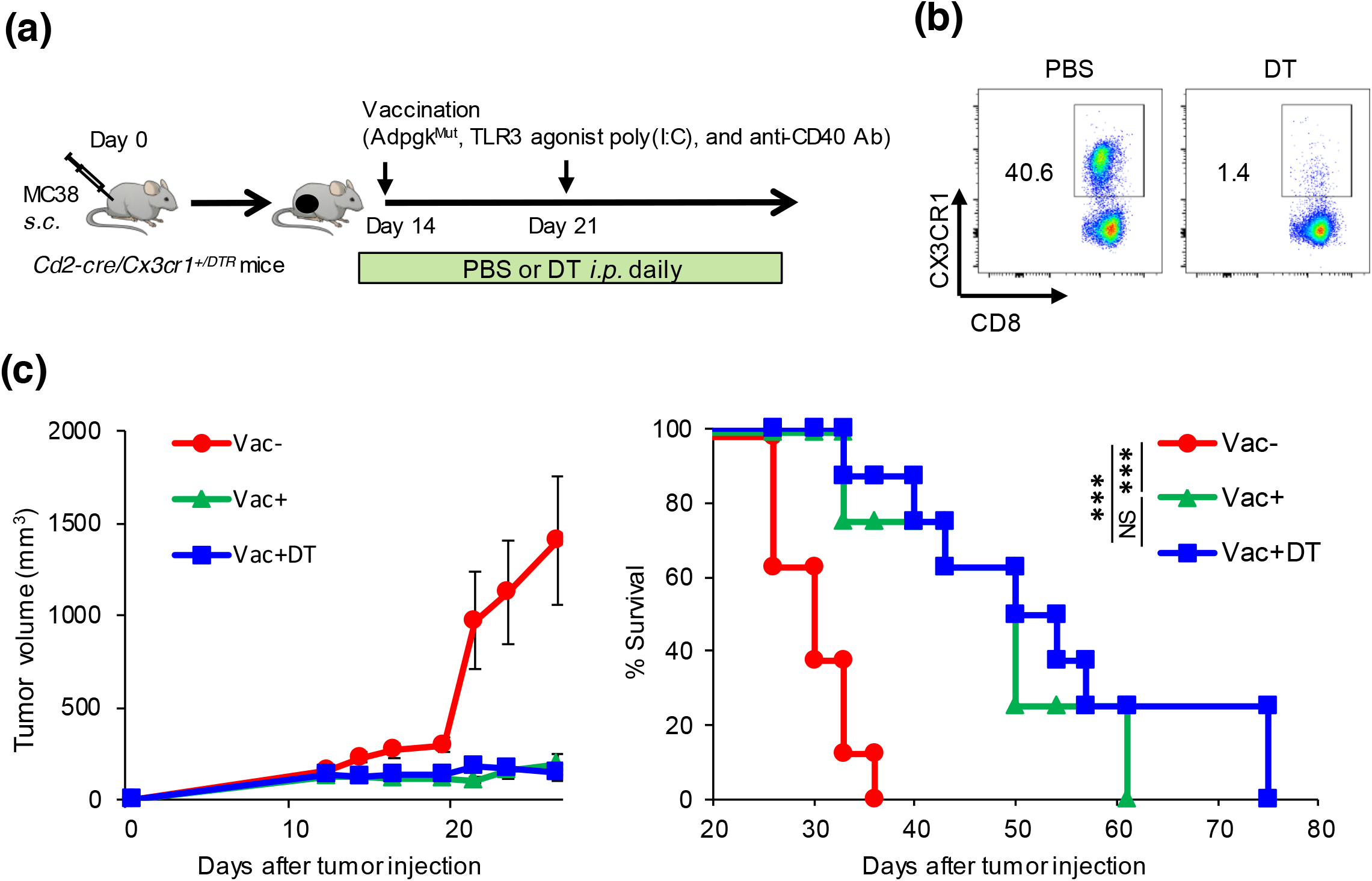
*In vivo* antitumor efficacy of antigen-specific CX3CR1 ^+^CD8^+^ T cells elicited by neoantigen/TLR3/CD40 agonists vaccination. (**a**) Schematic illustration showing evaluation of *in vivo* antitumor efficacy of the CX3CR1^+^ subset using MC38 tumor-bearing *Cd2-cre/Cx3cr1^+/DTR^* mice treated with Adpgk^Mut^/TLR3/CD40 agonists vaccination (Vac) followed by diphtheria toxin (DT) administration. DT (250 ng) was injected intraperitoneally every day from day 13. (**b**) Representative FACS plots showing selective depletion of CX3CR1^+^ subset in PB upon DT injection in *Cd2-cre/Cx3cr1*^+/*DTR*^ mice treated with Vac. Expression of CX3CR1 in CD8^+^ T cells in peripheral blood of mice treated with PBS or DT are shown. Numbers denote percentage of the CX3CR1^+^ subset. (**c**) Tumor growth and survival curves of MC38 tumor-bearing *Cd2-cre/Cx3cr1*^+/*DTR*^ mice in different treatment groups. (n ≥ 7 mice per group). Data shown are representative of two independent experiments. Data shown are representative from two independent experiments. \*\*\**P* < 0.001, \*\*\*\**P* < 0.0001 by log-rank (Mantel-Cox) test (**c**) or paired two-tailed *t*-test (**d**). Mean ± SEM.

### CD40 signaling is required for antitumor efficacy of neoantigen/TLR3/CD40 agonists and generation of antigen-specific CX3CR1^+^ CD8^+^ T cells

Neoantigen vaccination with a TLR3 agonist increases antigen-specific tumor-infiltrating lymphocytes in a preclinical model (31) and patients (4, 5). CD40 signaling drives robust DC activation to generate cytotoxic CD8^+^ T cells (32, 33), and synergizes with the TLR3 ligand in inducing an expansion of peptide or DC vaccine-primed and adoptively transferred antigen-specific CD8^+^ T cells (14–17). To delineate the role of CD40 signaling in the context of neoantigen vaccination, we examined antitumor efficacy of Adpgk^Mut^/TLR3/CD40 vaccination and generation of Adpgk^Mut^-specific CX3CR1^+^ CD8^+^ T cells using CD40^−/−^ mice. Antitumor efficacy of Adpgk^Mut^**/**TLR3/CD40 vaccination was abrogated in CD40^−/−^ mice (**Figure 4A**). The lack of increase in the frequency of Adpgk^Mut^-specific CX3CR1^+^ CD8^+^ T cells in CD40^−/−^ mice denoted the requirement of CD40-CD40 ligand pathway in this process (**Figure 4B**), in line with recent studies showing that CD4^+^ T-cell help or agonistic anti-CD40 Ab facilitates generation of CX3CR1^+^ CD8^+^ T cells (24, 34).

**Figure 4.**
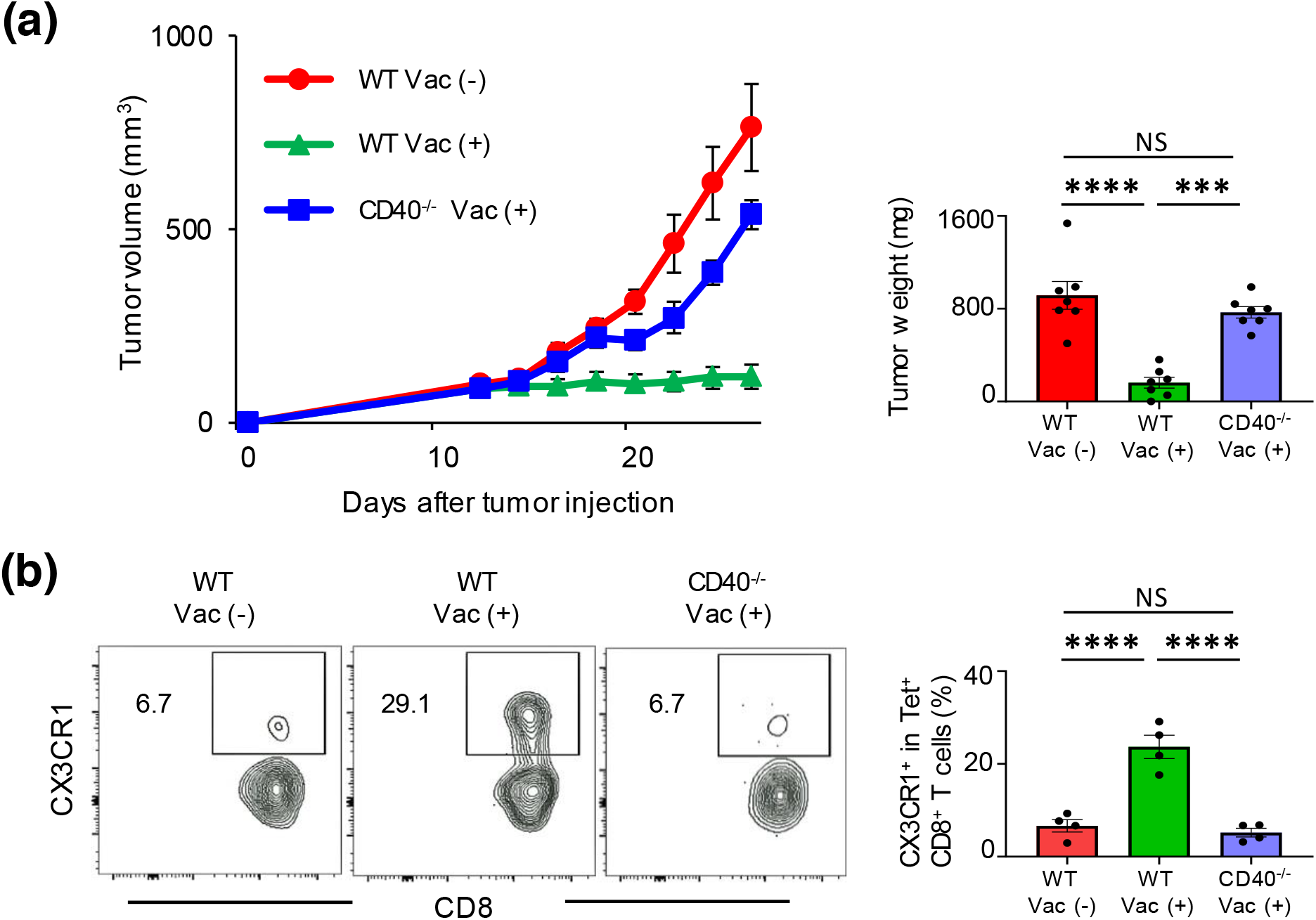
CD40-dependent generation of antigen-specific CX3CR1^+^CD8^+^ T-cell subsets upon neoantigen/TLR3/CD40 agonists vaccination. (**a**) Left shows tumor volume curves in MC38 tumor-bearing wild type C57BL/6 mice (WT) or CD40^−/−^ mice treated with or without Adpgk^Mut^/TLR3/CD40 agonists vaccination (Vac) (n = 7 mice per group). Right shows tumor weight. Tumors were harvested 1 week after second vaccination (Vac). (**b**) Left shows representative FACS plots showing CD8 and CX3CR1 expression gated with Adpgk^Mut^ Tet^+^CD8^+^ T cells in peripheral blood (PB) of MC38-tumor bearing WT or CD40 ^−/−^ mice treated with or without Vac. Numbers denote percentage of the CX3CR1^+^ subset. Right shows the frequency of the CX3CR1^+^ subset in each group (n = 4 mice per group). PB was harvested 1 week after second Vac. Data shown are representative of two independent experiments. NS: not significant, \*\*\**P* <0.001, \*\*\*\**P* <0.0001 by one-way ANOVA with Tukey’s multiple comparisons. Mean ± SEM.

### A Critical role of CD40 signaling in cDC1s for antitumor efficacy of neoantigen/TLR3/CD40 agonists vaccination and generation of antigen-specific CX3CR1^+^ CD8^+^ T cells

Batf3-dependent conventional type 1 dendritic cells (cDC1s) specialize in cross-presentation of exogenous antigens on MHC class I molecules, and play a key role in antitumor immunity and response to immunotherapy (14, 18, 31, 35–37). To assess whether cDC1s are necessary for the generation of CX3CR1^+^ CD8^+^ T cells, we employed mice deficient in Batf3 transcription factor and lack cDC1s (35). We found no therapeutic efficacy of Adpgk^Mut^**/**TLR3/CD40 vaccination was observed in Batf3^−/−^ mice (**Figure 5A**) along with a significant reduction of CX3CR1^+^ Tet^+^ CD8^+^ T cells (**Figure 5B**), suggesting the requirement of cDC1s for generation of antigen-specific CX3CR1^+^ CD8^+^ T cells.

**Figure 5.**
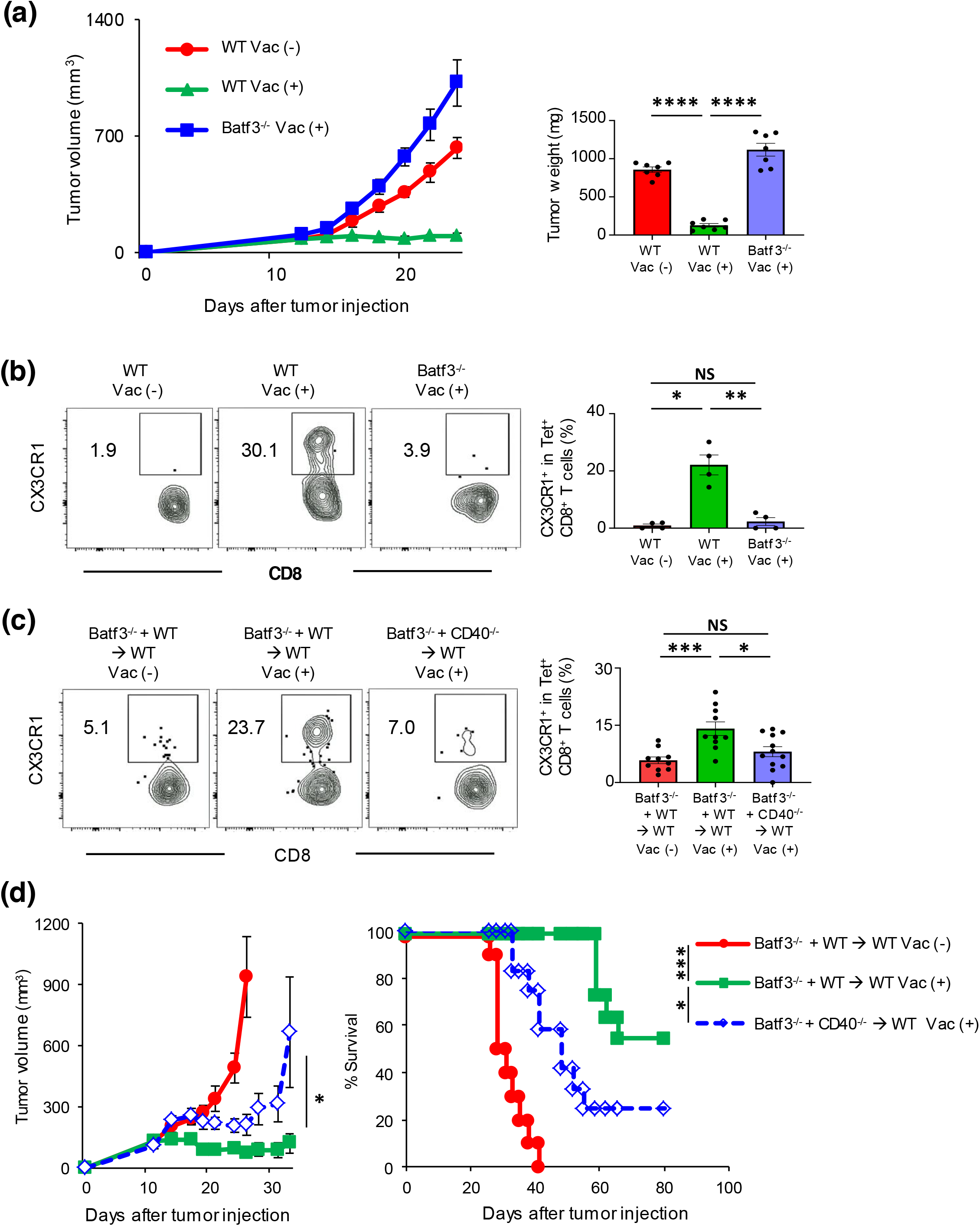
Critical roles of CD40 signaling in cDC1s for antitumor efficacy of neoantigen/TLR3/CD40 agonists vaccination and generation of antigen-specific CX3CR1^+^CD8^+^ T cells. (**a**) Left shows tumor volume curves in MC38 tumor-bearing wild type C57BL/6 mice (WT) or Batf3^−/−^ mice treated with or without Adpgk^Mut^/TLR3/CD40 agonists vaccination (Vac) (n = 7 mice per group). Tumors were harvested 1 week after second Vac, and tumor weight (mg) is shown in the right panel. (**b**) Left shows representative FACS plots showing CD8 and CX3CR1 expression gated with PB Adpgk^Mut^ Tet^+^CD8^+^ T cells in MC38 tumor-bearing WT or Batf3 ^−/−^ mice treated with or without Vac. Numbers denote percentage of the CX3CR1^+^ subset. Right shows the frequency of the CX3CR1^+^ subset in each group (n = 4 mice per group). (**c**) Left shows representative FACS plots showing CD8 and CX3CR1 expression gated with Adpgk^Mut^ Tet^+^ CD8^+^ T cells in PB of MC38-tumor bearing bone marrow chimera mice (Batf3^−/−^ and WT→WT or Batf3^−/−^ and CD40^−/−^→WT mice) treated with or without Vac. PB was harvested 1 week after second Vac (**b** and **c**). Data shown are representative of two independent experiments (**a**, **b** and **c**). (**d**) Tumor growth and survival curves in MC38 tumor-bearing mixed bone marrow chimera mice (Batf3^−/−^ and WT→WT or Batf3^−/−^ and CD40^−/−^→WT mice) treated with or without Vac (n = 10-11 mice per group). Data shown are pooled from two independent experiments. \**P* < 0.05, \*\**P* < 0.01, \*\*\**P* < 0.001, \*\*\*\**P* < 0.0001 by one-way ANOVA with Tukey’s multiple comparisons (**a**, **b**and **c**), Mann-Whitney *U* test (**d** for tumor volume), or log-rank (Mantel-Cox) test (d for survival). Mean ± SEM.

cDC1s express high levels of TLR3, and a TLR3 agonist poly(I:C) has been used in personalized neoantigen vaccine trials with vaccine-related objective responses (5–7). However, the role of CD40 signaling in cDC1s in vaccine-based immunotherapy remains elusive. To address a direct role of CD40 signaling in cDC1 lineage for CX3CR1^+^ CD8^+^ T cell development, we generated CD40^−/−^/Batf3^−/−^ or wild type (WT)/Batf3^−/−^ mixed bone marrow chimeras into irradiated C57BL/6 WT mice. CD40^−/−^/Batf3^−/−^ bone marrow transplanted recipient mice (CD40^−/−^ plus Batf3^−/−^ mixed bone marrow chimeras) lack CD40 expression only in Batf3-dependent bone marrow-derived cells while WT/Batf3^−/−^ bone marrow transplanted recipient mice have an intact CD40 signaling. In CD40^−/−^ plus Batf3^−/−^ mixed bone marrow chimeras, CX3CR1^+^ CD8^+^ T-cell responses remained minimal compared to WT plus Batf3^−/−^ mixed bone marrow chimeras, indicating that CD40 expression on cDC1s is pivotal for the CX3CR1^+^ CD8^+^ T-cell induction (**Figure 5C**). In line with these results, mice reconstituted with CD40^−/−^ plus Batf3^−/−^ mixed bone marrow chimeras had substantially decreased antitumor efficacy of Adpgk^Mut^/TLR3/CD40 vaccination compared to mice with WT plus Batf3^−/−^ mixed bone marrow chimeras (**Figure 5D**).

### Non-redundant requirement of CD80/86 but not CD70 signaling in cDC1s for antitumor efficacy of neoantigen/TLR3/CD40 agonists vaccination and generation of antigen-specific CX3CR1^+^ CD8^+^ T cells

In addition to CD40, both CD70 and CD80/86 have been reported to play an important role in DC-mediated T cell activation and persistence (38); however, the role of these molecules on cDC1s for neoantigen-specific T-cell differentiation remains elusive. Therefore, the requirements of CD70 and CD80/86 signaling in cDC1s were assessed in mixed bone marrow chimeric mice of CD70^−/−^ plus Batf3^−/−^ and CD80/86^−/−^ plus Batf3^−/−^. There was no difference in the frequency of Adpgk^Mut^-specific CX3CR1^+^ CD8^+^ T cells between WT plus Batf3^−/−^ and CD70^−/−^ plus Batf3^−/−^ mixed bone marrow chimeras (**Figure 6A**) with similar tumor control (**Figure 6B**), suggesting a redundant role of CD70 signaling in cDC1s. In sharp contrast, generation of Adpgk^Mut^-specific CX3CR1^+^ CD8^+^ T cells (**Figure 6C**) and antitumor efficacy of Adpgk^Mut^/TLR3/CD40 vaccination (**Figure 6D**) were abrogated in the CD80/86^−/−^ plus Batf3^−/−^ compared to the control WT plus Batf3^−/−^ mixed bone marrow chimeras. Thus, the CD80/86-CD28 pathway in cDC1s plays a crucial role in mediating antitumor efficacy and generating antigen-specific effector CX3CR1^+^ CD8^+^ T cells in the context of neoantigen vaccination. Taken together, effective neoantigen/TLR3/CD40 agonist vaccination correlates with generation and frequency of circulating antigen-specific CX3CR1^+^ CD8^+^ T cells mediated by CD40 and CD80/86 signaling in cDC1s.

**Figure 6.**
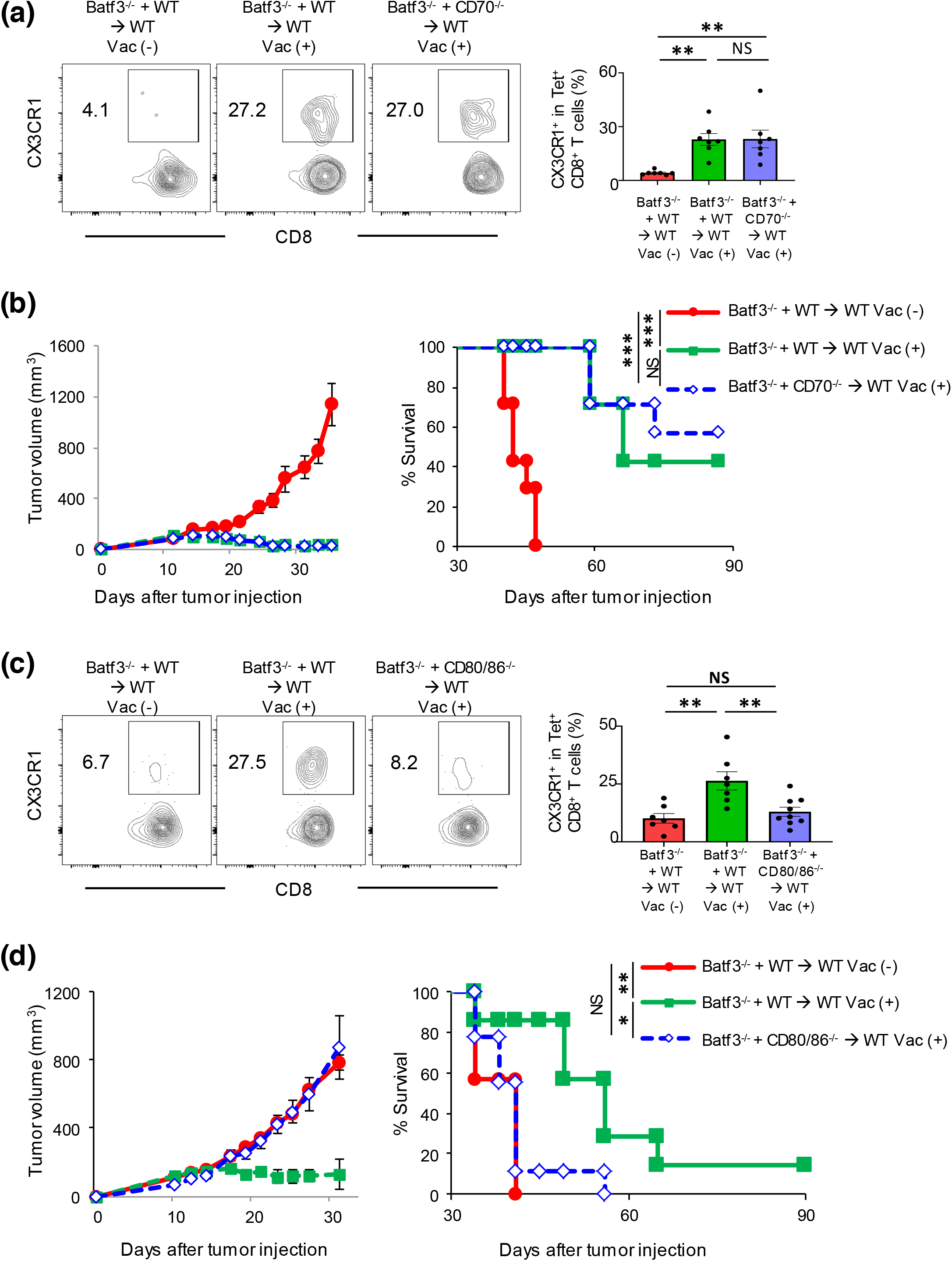
Roles of CD80/86 and CD70 signaling in cDC1s for antitumor efficacy of neoantigen/TLR3/CD40 agonists vaccination and generation of antigen-specific CX3CR1^+^CD8^+^ T cells. (**a**) Left shows representative FACS plots showing CD8 and CX3CR1 expression gated with Adpgk^Mut^ Tet^+^ CD8^+^ T cells in PB of MC38-tumor bearing bone marrow chimera mice (Batf3^−/−^ and WT→WT or Batf3^−/−^ and CD70^−/−^→WT mice) treated with or without Adpgk^Mut^/TLR3/CD40 agonists vaccination (Vac) Right shows the frequency of the CX3CR1^+^Tet^+^CD8^+^ T cells in each group (n = 7 mice per group). (**b**) Tumor growth and survival curves of MC38 tumor-bearing mixed bone marrow chimera mice in different treatment groups as indicated. (n = 7 mice per group) (**c**) Left shows representative FACS plots showing CD8 and CX3CR1 expression gated with Adpgk^Mut^ Tet^+^ CD8^+^ T cells in PB of MC38-tumor bearing bone marrow chimera mice (Batf3^−/−^ and WT→WT or Batf3^−/−^ and CD80/86^−/−^→WT mice) treated with or without Vac (n = 7-9 mice per group). (**d**) Tumor growth and survival curves of MC38 tumor-bearing mixed bone marrow chimera mice in different treatment groups as indicated (n = 7-9 mice per group). PB was harvested 1 week after second Vac (**a** and **c**). Data shown are representative from two or three independent experiments. NS not significant, \**P* <0.05, \*\**P* <0.01, \*\*\**P* <0.001 by one-way ANOVA with Tukey’s multiple comparisons (**a**, **c**) or log-rank (Mantel-Cox) test (**b**, **d**). Mean ± SEM.

## Discussion

In this study we have shown potent antitumor efficacy of neoantigen/TLR3/CD40 stimulations in a preclinical model, and identified a novel link between antitumor efficacy of neoantigen vaccine and an increased frequency of circulating neoantigen-specific CX3CR1^+^ CD8^+^ T cells. Moreover, using mixed bone marrow chimeras from Batf3^−/−^, CD40^−/−^, CD70^−/−^ and CD80/86^−/−^ mice, we have revealed a pivotal role of CD40 and CD80/86 signaling in cDC1s for generation of neoantigen-specific CX3CR1^+^ CD8^+^ T cells and *in vivo* antitumor efficacy of neoantigen/TLR3/CD40 agonists vaccination.

Engagement of CD40 on DCs induces costimulatory molecules on their surface, promotes their cytokine production, facilitates the cross-presentation of antigen (32), and synergistically augments expansion of both endogenous and adoptive transferred T cells with TLR agonist (14–19). Our data further demonstrated that among the diversity of the DC subsets cDC1s play a critical role in antitumor efficacy of TLR3/CD40 stimulation. Although poly(I:C) (TLR3 agonist) has been used in neoantigen vaccine clinical studies (5, 6), these findings underscore the implication of agonistic anti-CD40 Ab in combination with poly(I:C) for the maximal engagement of cDC1 in vaccine-based therapy.

The molecular and cellular mechanisms underlying differentiation of CX3CR1^−^ to CX3CR1^+^ CD8^+^ T cells remain elusive. Recent studies have indicated that CD4^+^ T-cell help or agonistic anti-CD40 Ab facilitates generation of CX3CR1^+^ CD8^+^ T cells (24, 34). Our results further demonstrated the non-redundant requirement of CD40 signaling in cDC1s for the generation of antigen-specific CX3CR1^+^ CD8^+^ T cells. The observation that CD70 signaling in cDC1s was dispensable for generation of the CX3CR1^+^ subset was unexpected given the several lines of evidence suggesting that CD40 signaling relays the CD4^+^ T-cell help signal by the induction of CD70 expression on DCs (14, 39). However, not all molecules regulated by CD4^+^ T-cell help are mediated by CD27, and cytokine signaling may have contributed to the generation of the CX3CR1^+^ subset (34). In support of this notion, IL-21 was found to modulate T-cell differentiation and generation of CX3CR1^+^ CD8^+^ T cells (24, 40). Additional mechanistic studies are needed to further determine the signaling required for generation of the CX3CR1^+^ subset.

The lack of a reliable surrogate marker for clinical response remains the major limitation of vaccine clinical trials (1, 2). Mere presence of vaccine-primed tumor-specific T cells might not correlate with effective regression of the tumor, and cannot be used as a standalone biomarker for vaccine efficacy (41). In our study, although frequency of PB antigen-specific total CD8^+^ T cells had a positive inverse correlation with tumor volume, differentiation of antigen-specific CD8^+^ T cells defined by CX3CR1 expression had a markedly higher inverse correlation with tumor volume. Furthermore, our findings of an increased frequency of CX3CR1^+^ CD8^+^ T cells that have the ability to produce IFN-γ, TNF-α and GZMA after successful vaccination align with the evidence from previous clinical studies showing that functional assessment of T cells such as IFN-γ production and/or cytolytic activity was correlated with response in patients treated with peptide vaccines (21–23).

An increased frequency of the CX3CR1^+^ subset was identified not only in Tet^+^ CD8^+^ T cells, but also in total CD8^+^ T cells, which had a strong inverse correlation with tumor volume. It remains unclear, however, whether this was due to an increased frequency of non-specific CX3CR1^+^ CD8^+^ T cells or an epitope-spreading phenomenon, the spread of the immune response from one antigen to another antigen expressed in the same tissue (1, 2). Additional studies are required to determine whether Adpgk^Mut^/TLR3/CD40 vaccination-induced CX3CR1^+^ CD8^+^ T cells contain tumor-specific T cells other than Adpgk^Mut^-specific T cells. Nevertheless, this observation suggests the potential utility of CX3CR1 as a blood-based predictive T-cell biomarker for other immunotherapies such as immune checkpoint inhibitor therapy. Indeed, an increased frequency of PB CX3CR1^+^ CD8^+^ T cells was observed in patients who responded to immune checkpoint inhibitors (28).

Our work and others have provided some insights into the theoretical advantages of CX3CR1 as a circulating surrogate marker for vaccine efficacy. Vaccine-primed CX3CR1^+^ CD8^+^ T cells expressed low levels of CD62L and CXCR3, trafficking receptors necessary for entry across lymphoid organ HEV and the tumor microvasculature, respectively (11, 12), that may have contributed to increase its frequency in PB. Furthermore, unlike other T-cell proliferation, activation or co-stimulatory/inhibitory molecules transiently expressed after activation, CX3CR1 is stably expressed on CD8^+^ T cells through unidirectional differentiation from CX3CR1^−^ CD8^+^ T cells during the effector phase (8–10).

Although the scope of our study is limited to CX3CR1, another future area of investigation would be to compare the utility of CX3CR1 as a circulating surrogate marker for vaccine efficacy with other molecules such as PD-1, which was reported to be expressed in circulating neoantigen-specific CD8^+^ T cells (42). Notably, our study was limited by the use of a transplantable non-orthotopic mouse tumor model with high mutational burden. Thus, the utility of CX3CR1 as a circulating T-cell biomarker for vaccine therapy targeting endogenous tumor-associated antigens or tumors with lower mutational burden remains uncertain. Further studies in genetically engineered mouse models or mouse tumors harboring endogenous tumor-associated antigen are warranted to better understand the overall utility of CX3CR1 in vaccine therapy.

Our *in vivo* depletion study revealed negligible contribution of CX3CR1^+^ CD8^+^ T cells elicited by Adpgk^Mut^/TLR3/CD40 vaccination against established MC38 tumors, in line with a previous study using a preclinical model of adoptive T cell therapy against B16 melanoma (10). These findings are in parallel to the compelling evidence that terminally-differentiated effector CD8^+^ T cells have less antitumor efficacy *in vivo* compared to less-differentiated T cells (43), but in contrast to previous studies demonstrating decreased antitumor immunity in CX3CR1^−/−^ mice or with the use of the CX3CR1 antagonist (25–28, 34). A possible explanation is that in *Cd2-cre/Cx3cr1^+/DTR^* mice, DT administration does not affect monocytes, DCs, M-MDSCs and TAMs that may express CX3CR1 (29, 30) and negatively regulate antitumor immunity. Another possibility is that CX3CR1/CX3CL1 pathway might not play a significant role in an MC38 tumor model, and trafficking of CX3CR1^+^ CD8^+^ T cells to the TME is not efficient in the setting of their decreased CXCR3 expression (11). Previous preclinical studies using mouse and xenograft tumor models revealed that augmenting CX3CR1/CX3CL1 axis enhances T cell/NK-cell mediated antitumor efficacy (25, 44, 45). Therefore, the role of CX3CR1^+^ CD8^+^ T cells in antitumor reactivity against solid malignancies might be context-dependent, and remains to be elucidated.

In summary, our study revealed cDC1-mediated potent antitumor efficacy of neoantigen vaccine with dual TLR3/CD40 stimulation, identified a correlation between antitumor response and the frequency of antigen-specific PB CX3CR1^+^ CD8^+^ T cells, and provided the mechanistic insight into the generation of CX3CR1^+^ CD8^+^ T cells. These data suggest the maximal engagement of cDC1s with TLR3/CD40 stimulation for the development of more effective neoantigen vaccine therapy, and the potential implications of CX3CR1 as a circulating T-cell predictive biomarker for response in future clinical studies.

## Abbreviations

Ab: Antibody
Batf3: Basic leucine zipper transcription factor ATF-like 3
cDC1: Conventional type 1 dendritic cell
CTLA-4: Cytotoxic T-lymphocyte-associated protein 4
CX3CR1: CX3C chemokine receptor 1
DC: Dendritic cell
DT: Diphtheria toxin
GZMA: Granzyme A
HEV: High endothelial venules
ICOS: Inducible T cell co-stimulator
IFN-γ: Interferon gamma
KLRG1: Killer-cell lectin like receptor G1
MHC: Major histocompatibility complex
NT: No treatment
PB: Peripheral blood
PD-1: Programmed cell death protein 1
poly(I:C): Polyinosinic-polycytidylic acid sodium salt
TAM: Tumor associated macrophage
TLR3: Toll-like receptor 3
TME: Tumor microenvironment
TNF-α: Tumor necrosis factor alpha
WT: Wild-type

## Acknowledgments

We acknowledge the NIH Tetramer Core Facility (contract HHSN272201300006C) for provision of MHC class I tetramers, Dr. Weiping Zou (University of Michigan, Ann Arbor, Michigan) for MC38 cells, Dr. James J. Moon (University of Michigan, Ann Arbor, Michigan) and Dr. Toshihiro Yokoi (Roswell Park) for technical assistance, and Dr. Suzanne M. Hess and Ms. Judith Epstein for administrative assistance. This work was supported by National Cancer Institute (NCI) grant P30CA016056 involving the use of Roswell Park’s Flow and Image Cytometry, Genomic Shared, and Biostatistics & Statistical Genomics Resources. This work was supported by Roswell Park Alliance Foundation and NCI/ K08CA197966 (F. Ito), Uehara Memorial Foundation (T. Oba), and Astellas Foundation for Research on Metabolic Disorders and the Nakatomi Foundation (T. Yamauchi).

## Declarations

### Ethics approval

All animal studies were reviewed and approved by the Roswell Park institutional animal care and use program and facilities (protocol #1316M and 1356M). All aspects of animal research and husbandry were conducted in accordance with the federal Animal Welfare Act and the NIH Guide for the Care and Use of Laboratory Animals.

### Consent for publication

All author consent for the publication.

### Availability of data and material

All data generated or analyzed during this study are included in this published article and supplementary information file.

### Author contributions

T.Y. contributed development of methodology, acquisition of data, analysis and interpretation of data, writing, review, and revision of the manuscript. T.H, contributed development of methodology, acquisition of data, analysis and interpretation of data, review, and revision of the manuscript. T.O. contributed acquisition of data, analysis and interpretation of data, review, and revision of the manuscript. K.A. contributed analysis and interpretation of data, review, and revision of the manuscript. X.C. provided CD70^−/−^ mice, review, and revision of the manuscript. F.I. developed the concept, managed the project, coordinated author activities, and provided final approval of the version to be submitted.

---

**Supplementary Figure 1.**
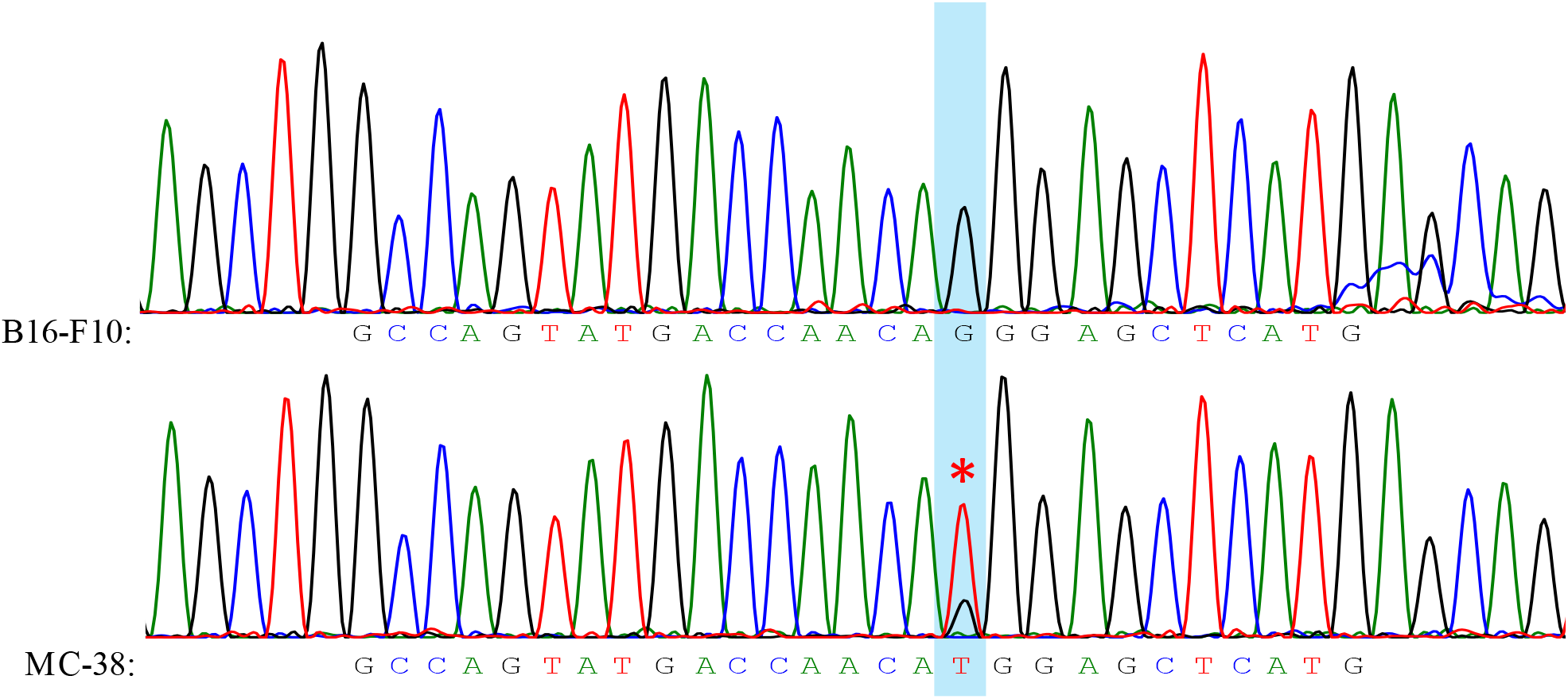
cDNA sequencing of mutated Adpgk neoantigen in MC38 cells. Total RNA was extracted from MC38 cells (harboring mutated Adpgk) along with B16F10 cells (harboring wild type Adpgk), and subjected to cDNA synthesis for Sanger DNA sequencing. An asterisk indicates the mutation of G→T, resulting in arginine (R) to methionine (M) transformation on 304th amino acid sequences of Adpgk gene.

**Supplementary Figure 2.**
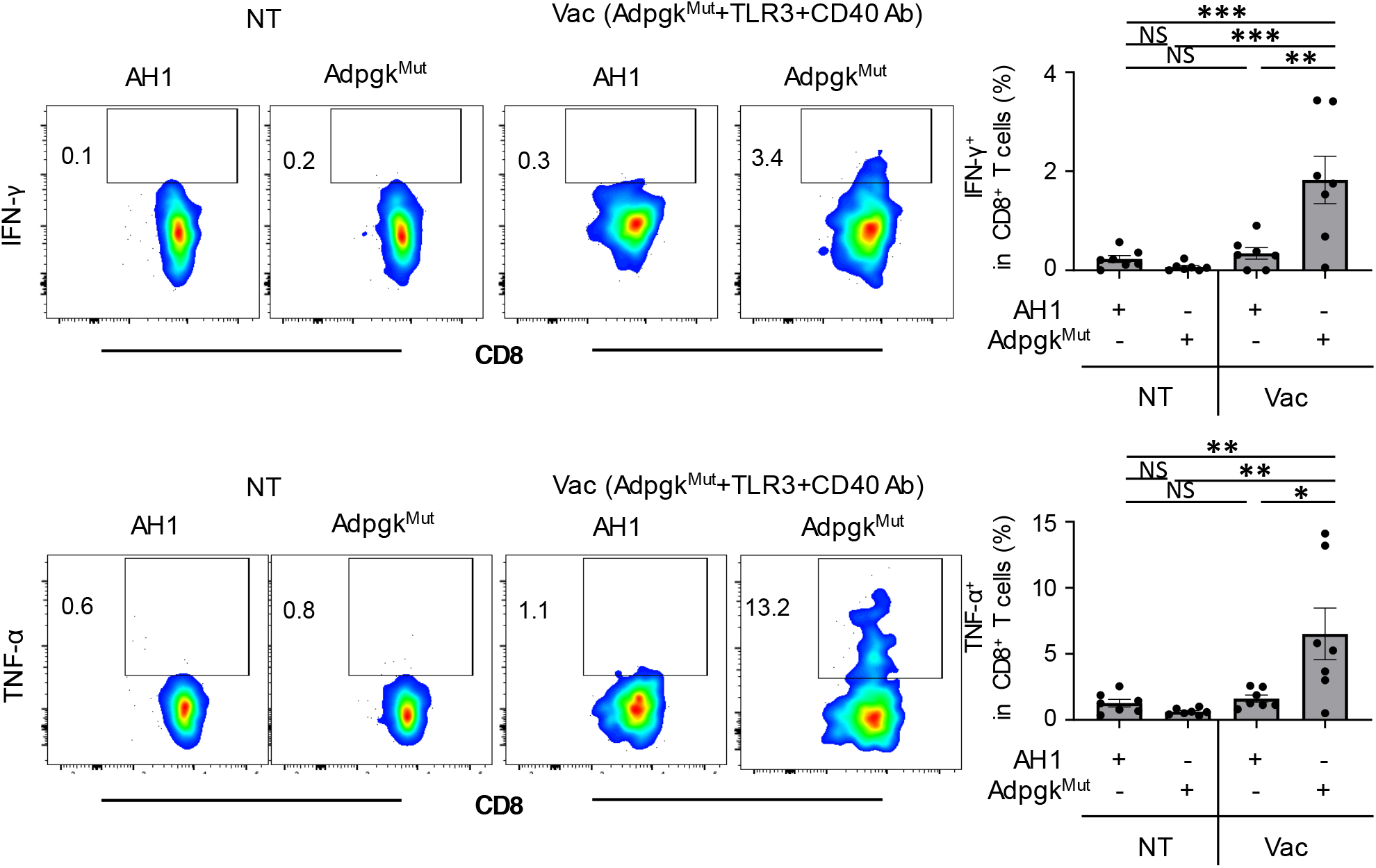
Neoantigen/TLR3/CD40 vaccination facilitates neoantigen-specific CD8^+^ T cell infiltration into tumors. Left shows representative FACS plots showing CD8 and IFN-γ (upper) or TNF-α (lower) expression gated with CD8^+^ T cells in tumors of MC38-tumor bearing mice treated with PBS (NT) or neoantigen/TLR3/CD40 vaccinations. Tumor cells were co-cultured with AH1 or Adpgk peptide in the presence of blefeldin A for 5 hrs before intracellular staining. Numbers denote percentage of IFN-γ^+^ or TNF-α^+^ cells. Right panel shows the frequency of the IFN-γ^+^ cells in CD8^+^ T cells in each group (n = 7 mice per group). Data shown are representative from two or three independent experiments. NS not significant, \**P* < 0.05, \*\**P* < 0.01, \*\*\**P* < 0.001 by one-way ANOVA with Tukey’s multiple comparisons. Mean ± SEM.

**Supplementary Figure 3.**
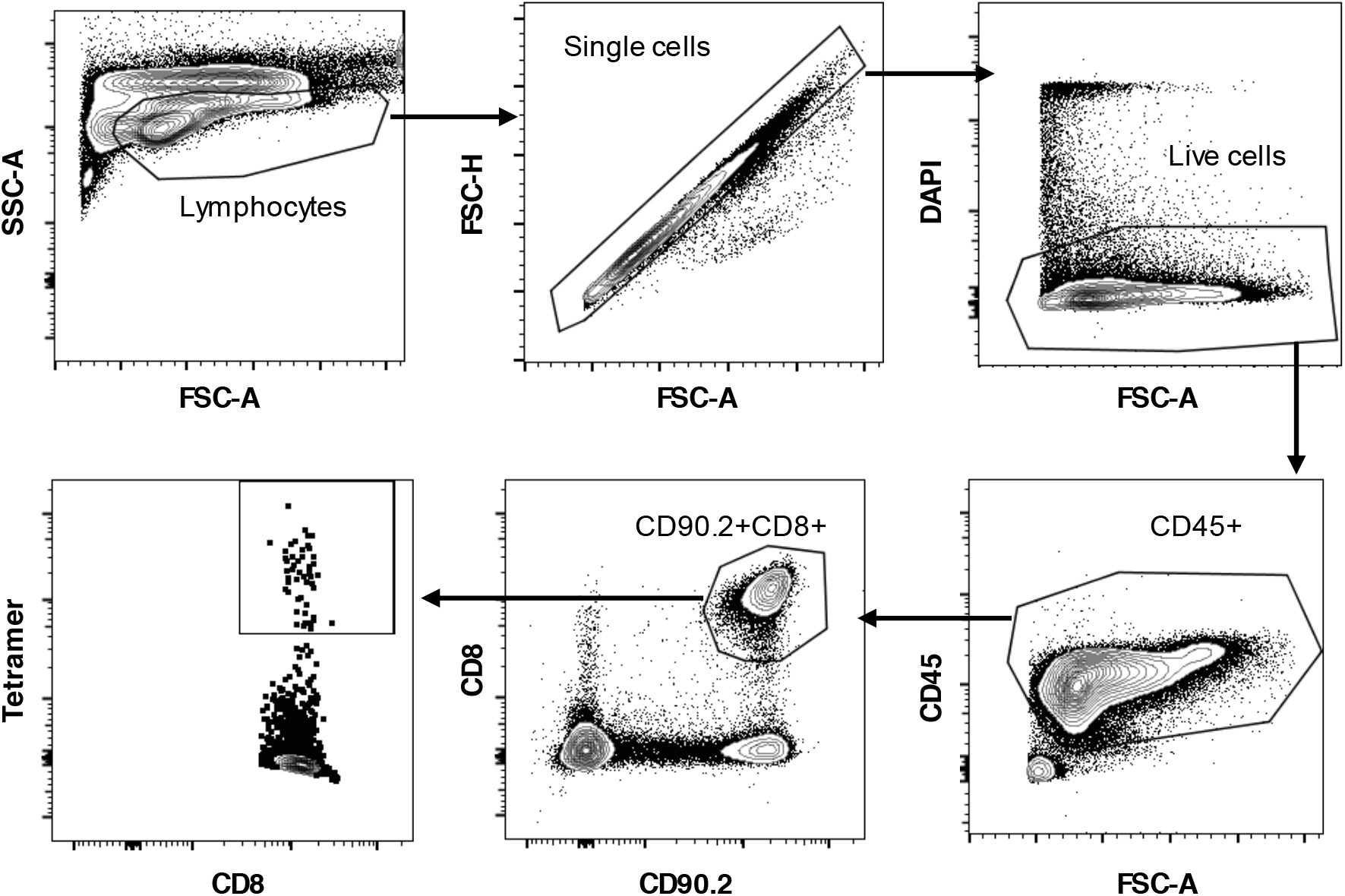
Flow cytometry and gating of circulating Adpgk^Mut^ eoantigen-specific CD8^+^ T cells in MC38-tumor bearing mice. Representative of greater than 5 independent experiments.

**Supplementary Figure 4.**
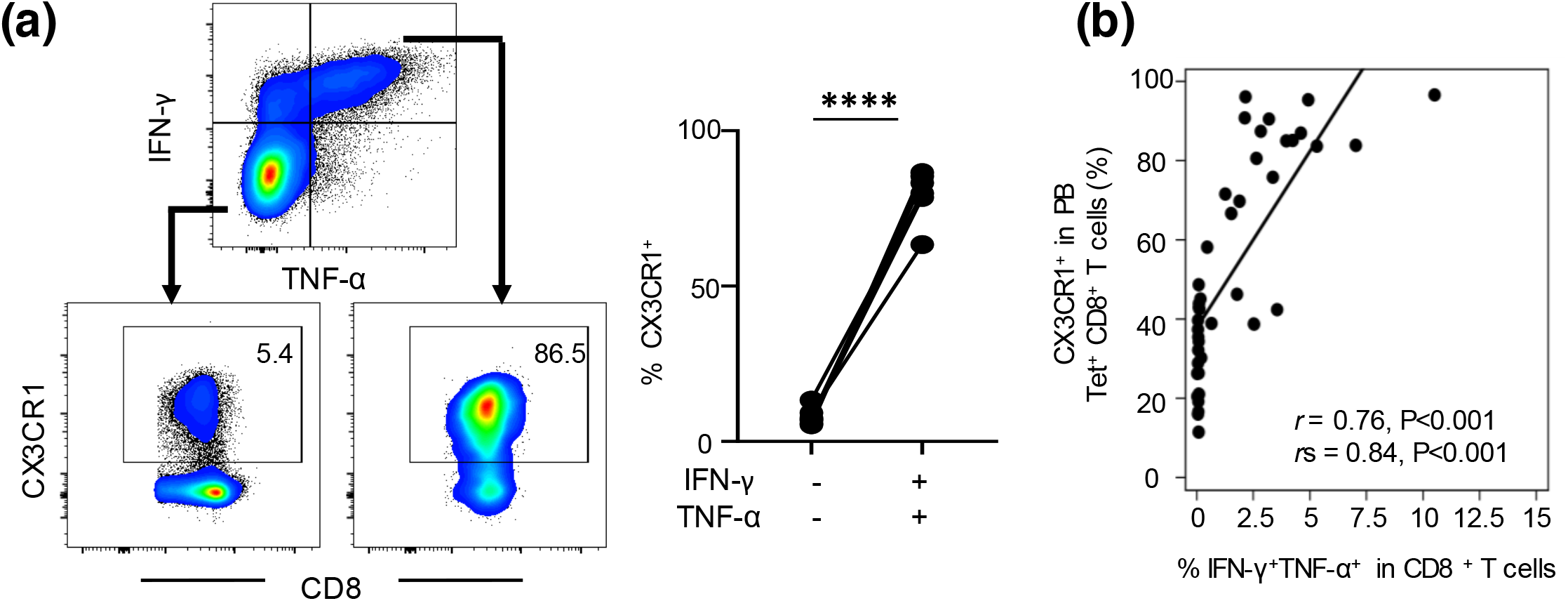
High frequency of CX3CR1^+^ subset in neoantigen/TLR3/CD40 vaccination-induced peripheral CD8^+^ T cells that are capable of producing effector cytokines. (**a**) Representative FACS plots showing IFN-γ and TNF-α expression gated with CD8^+^ T cells in splenocytes of MC38-tumor bearing mice treated with Adpgk^Mut^/TLR3/CD40 vaccinations. Lower panels show CX3CR1 expression in IFN-γ^−^ TNF-α^−^ cells (left) and IFN-γ^+^ TNFF-α^+^ cells (right). Numbers denote percentage of CX3CR1^+^ cells. Data shown are representative of three independent experiments. Frequency of CX3CR1 cells from the IFN-γ^−^ TNF-α^−^ or IFN-γ^+^ TNF-α^+^ subset is shown in the right panel (n = 7 mice per group). Spleens were harvested 1 week after second vaccination. \*\*\*\**P* < 0.0001 by paired two-tailed *t*-test **(a**). (**b**) Scatter plot of the frequency of IFN-γ^+^TNF-α^+^ cells against the frequency of Adpgk^Mut^ mer (Tet)^+^ CX3CR1^+^CD8^+^ T cells in splenocytes. Correlation is shown using Pearson lation (*r*) and Spearman correlation coefficients (*r*_s_). Data shown are representative of two independent experiments.

**Table S1:**
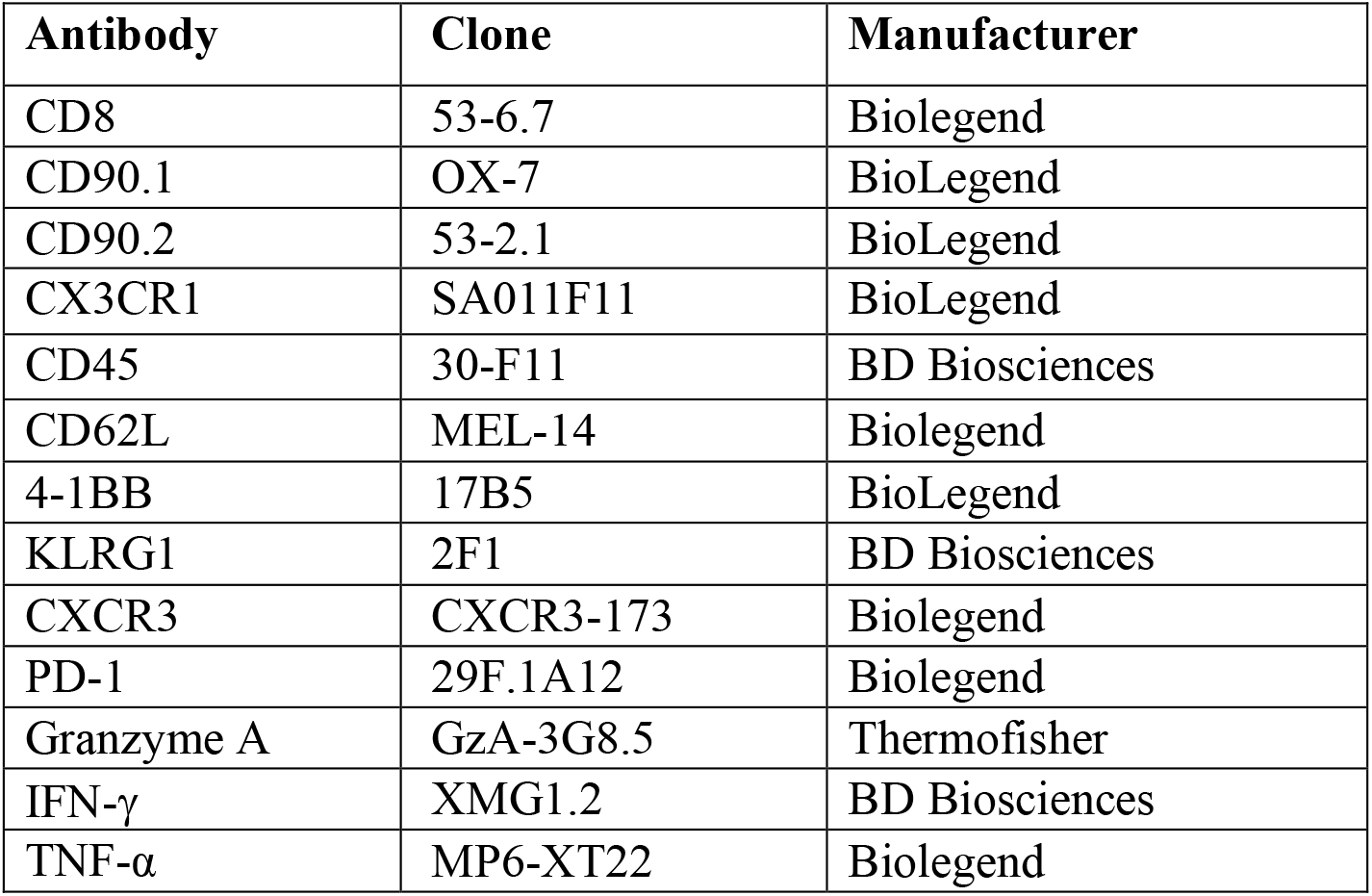
Fluorochrome-conjugated antibodies used in flow cytometry

## References

1. Butterfield LH (2015) Cancer vaccines. BMJ (Clinical research ed.). 350: h988. doi: 10.1136/bmj.h988

2. Melero I, Gaudernack G, Gerritsen W et al. (2014) Therapeutic vaccines for cancer: an overview of clinical trials. Nature reviews. Clinical oncology. 11: 509–24. doi: 10.1038/nrclinonc.2014.111

3. Schumacher TN, Schreiber RD (2015) Neoantigens in cancer immunotherapy. Science. 348: 69–74. doi: 10.1126/science.aaa4971

4. Sahin U, Derhovanessian E, Miller M et al. (2017) Personalized RNA mutanome vaccines mobilize poly-specific therapeutic immunity against cancer. Nature. 547: 222–6. doi: 10.1038/nature23003

5. Keskin DB, Anandappa AJ, Sun J et al. (2019) Neoantigen vaccine generates intratumoral T cell responses in phase Ib glioblastoma trial. Nature. 565: 234–9. doi: 10.1038/s41586-018-0792-9

6. Ott PA, Hu Z, Keskin DB et al. (2017) An immunogenic personal neoantigen vaccine for patients with melanoma. Nature. 547: 217–21. doi: 10.1038/nature22991

7. Hilf N, Kuttruff-Coqui S, Frenzel K et al. (2019) Actively personalized vaccination trial for newly diagnosed glioblastoma. Nature. 565: 240–5. doi: 10.1038/s41586-018-0810-y

8. Gordon CL, Lee LN, Swadling L et al. (2018) Induction and Maintenance of CX3CR1-Intermediate Peripheral Memory CD8(+) T Cells by Persistent Viruses and Vaccines. Cell Rep. 23: 768–82. doi: 10.1016/j.celrep.2018.03.074

9. Gerlach C, Moseman EA, Loughhead SM, Alvarez D, Zwijnenburg AJ, Waanders L, Garg R, de la Torre JC, von Andrian UH (2016) The Chemokine Receptor CX3CR1 Defines Three Antigen-Experienced CD8 T Cell Subsets with Distinct Roles in Immune Surveillance and Homeostasis. Immunity. 45: 1270–84. doi: 10.1016/j.immuni.2016.10.018

10. Yamauchi T, Hoki T, Oba T, Saito H, Attwood K, Sabel MS, Chang AE, Odunsi K, Ito F (2020) CX3CR1-CD8+ T cells are critical in antitumor efficacy, but functionally suppressed in the tumor microenvironment. JCI insight. 5: e133920. doi: 10.1172/jci.insight.133920

11. Mikucki ME, Fisher DT, Matsuzaki J et al. (2015) Non-redundant requirement for CXCR3 signalling during tumoricidal T-cell trafficking across tumour vascular checkpoints. Nature communications. 6: 7458. doi: 10.1038/ncomms8458

12. von Andrian UH, Mempel TR (2003) Homing and cellular traffic in lymph nodes. Nat Rev Immunol. 3: 867–78. doi: 10.1038/nri1222

13. Yadav M, Jhunjhunwala S, Phung QT et al. (2014) Predicting immunogenic tumour mutations by combining mass spectrometry and exome sequencing. Nature. 515: 572–6. doi: 10.1038/nature14001

14. Oba T, Hoki T, Yamauchi T, Keler T, Marsh HC, Cao X, Ito F (2020) A Critical Role of CD40 and CD70 Signaling in Conventional Type 1 Dendritic Cells in Expansion and Antitumor Efficacy of Adoptively Transferred Tumor-Specific T Cells. J Immunol. 205: 1867–77. doi: 10.4049/jimmunol.2000347

15. Nimanong S, Ostroumov D, Wingerath J et al. (2017) CD40 Signaling Drives Potent Cellular Immune Responses in Heterologous Cancer Vaccinations. Cancer research. 77: 1918–26. doi: 10.1158/0008-5472.Can-16-2089

16. Cho HI, Celis E (2009) Optimized peptide vaccines eliciting extensive CD8 T-cell responses with therapeutic antitumor effects. Cancer research. 69: 9012–9. doi: 10.1158/0008-5472.Can-09-2019

17. Ahonen CL, Doxsee CL, McGurran SM, Riter TR, Wade WF, Barth RJ, Vasilakos JP, Noelle RJ, Kedl RM (2004) Combined TLR and CD40 triggering induces potent CD8+ T cell expansion with variable dependence on type I IFN. The Journal of experimental medicine. 199: 775–84. doi: 10.1084/jem.20031591

18. Oba T, Long MD, Keler T, Marsh HC, Minderman H, Abrams SI, Liu S, Ito F (2020) Overcoming primary and acquired resistance to anti-PD-L1 therapy by induction and activation of tumor-residing cDC1s. Nature communications. 11: 5415. doi: 10.1038/s41467-020-19192-z

19. Scarlett UK, Cubillos-Ruiz JR, Nesbeth YC, Martinez DG, Engle X, Gewirtz AT, Ahonen CL, Conejo-Garcia JR (2009) In situ stimulation of CD40 and Toll-like receptor 3 transforms ovarian cancer-infiltrating dendritic cells from immunosuppressive to immunostimulatory cells. Cancer research. 69: 7329–37. doi: 10.1158/0008-5472.Can-09-0835

20. Khalil DN, Suek N, Campesato LF et al. (2019) In situ vaccination with defined factors overcomes T cell exhaustion in distant tumors. J Clin Invest. 129: 3435–47. doi: 10.1172/jci128562

21. Carbone DP, Ciernik IF, Kelley MJ et al. (2005) Immunization with mutant p53- and K-ras-derived peptides in cancer patients: immune response and clinical outcome. J Clin Oncol. 23: 5099–107. doi: 10.1200/jco.2005.03.158

22. Kirkwood JM, Lee S, Moschos SJ, Albertini MR, Michalak JC, Sander C, Whiteside T, Butterfield LH, Weiner L (2009) Immunogenicity and antitumor effects of vaccination with peptide vaccine+/-granulocyte-monocyte colony-stimulating factor and/or IFN-alpha2b in advanced metastatic melanoma: Eastern Cooperative Oncology Group Phase II Trial E1696. Clin Cancer Res. 15: 1443–51. doi: 10.1158/1078-0432.Ccr-08-1231

23. Dillon PM, Olson WC, Czarkowski A, Petroni GR, Smolkin M, Grosh WW, Chianese-Bullock KA, Deacon DH, Slingluff CL, Jr. (2014) A melanoma helper peptide vaccine increases Th1 cytokine production by leukocytes in peripheral blood and immunized lymph nodes. Journal for immunotherapy of cancer. 2: 23. doi: 10.1186/2051-1426-2-23

24. Zander R, Schauder D, Xin G, Nguyen C, Wu X, Zajac A, Cui W (2019) CD4(+) T Cell Help Is Required for the Formation of a Cytolytic CD8(+) T Cell Subset that Protects against Chronic Infection and Cancer. Immunity. 51: 1028–42.e4. doi: 10.1016/j.immuni.2019.10.009

25. Xin H, Kikuchi T, Andarini S et al. (2005) Antitumor immune response by CX3CL1 fractalkine gene transfer depends on both NK and T cells. Eur J Immunol. 35: 1371–80. doi: 10.1002/eji.200526042

26. Zeng Y, Huebener N, Fest S et al. (2007) Fractalkine (CX3CL1)- and interleukin-2-enriched neuroblastoma microenvironment induces eradication of metastases mediated by T cells and natural killer cells. Cancer research. 67: 2331–8. doi: 10.1158/0008-5472.Can-06-3041

27. Yu YR, Fong AM, Combadiere C, Gao JL, Murphy PM, Patel DD (2007) Defective antitumor responses in CX3CR1-deficient mice. Int J Cancer. 121: 316–22. doi: 10.1002/ijc.22660

28. Yan Y, Cao S, Liu X et al. (2018) CX3CR1 identifies PD-1 therapy-responsive CD8+ T cells that withstand chemotherapy during cancer chemoimmunotherapy. JCI insight. 3: e97828. doi: 10.1172/jci.insight.97828

29. Nishimura M, Umehara H, Nakayama T, Yoneda O, Hieshima K, Kakizaki M, Dohmae N, Yoshie O, Imai T (2002) Dual functions of fractalkine/CX3C ligand 1 in trafficking of perforin+/granzyme B+ cytotoxic effector lymphocytes that are defined by CX3CR1 expression. J Immunol. 168: 6173–80.

30. Bronte V, Brandau S, Chen SH et al. (2016) Recommendations for myeloid-derived suppressor cell nomenclature and characterization standards. Nature communications. 7: 12150. doi: 10.1038/ncomms12150

31. Sanchez-Paulete AR, Cueto FJ, Martinez-Lopez M et al. (2016) Cancer Immunotherapy with Immunomodulatory Anti-CD137 and Anti-PD-1 Monoclonal Antibodies Requires BATF3-Dependent Dendritic Cells. Cancer Discov. 6: 71–9. doi: 10.1158/2159-8290.cd-15-0510

32. Elgueta R, Benson MJ, de Vries VC, Wasiuk A, Guo Y, Noelle RJ (2009) Molecular mechanism and function of CD40/CD40L engagement in the immune system. Immunol Rev. 229: 152–72. doi: 10.1111/j.1600-065X.2009.00782.x

33. Vonderheide RH (2018) The Immune Revolution: A Case for Priming, Not Checkpoint. Cancer Cell. 33: 563–9. doi: 10.1016/j.ccell.2018.03.008

34. Ahrends T, Spanjaard A, Pilzecker B, Babala N, Bovens A, Xiao Y, Jacobs H, Borst J (2017) CD4(+) T Cell Help Confers a Cytotoxic T Cell Effector Program Including Coinhibitory Receptor Downregulation and Increased Tissue Invasiveness. Immunity. 47: 848–61.e5. doi: 10.1016/j.immuni.2017.10.009

35. Hildner K, Edelson BT, Purtha WE et al. (2008) Batf3 deficiency reveals a critical role for CD8alpha+ dendritic cells in cytotoxic T cell immunity. Science. 322: 1097–100. doi: 10.1126/science.1164206

36. Broz ML, Binnewies M, Boldajipour B et al. (2014) Dissecting the tumor myeloid compartment reveals rare activating antigen-presenting cells critical for T cell immunity. Cancer Cell. 26: 638–52. doi: 10.1016/j.ccell.2014.09.007

37. Morrison AH, Diamond MS, Hay CA, Byrne KT, Vonderheide RH (2020) Sufficiency of CD40 activation and immune checkpoint blockade for T cell priming and tumor immunity. Proc Natl Acad Sci U S A. 117: 8022–31. doi: 10.1073/pnas.1918971117

38. Bak SP, Barnkob MS, Bai A, Higham EM, Wittrup KD, Chen J (2012) Differential requirement for CD70 and CD80/CD86 in dendritic cell-mediated activation of tumor-tolerized CD8 T cells. J Immunol. 189: 1708–16. doi: 10.4049/jimmunol.1201271

39. Borst J, Ahrends T, Babala N, Melief CJM, Kastenmuller W (2018) CD4(+) T cell help in cancer immunology and immunotherapy. Nat Rev Immunol. 18: 635–47. doi: 10.1038/s41577-018-0044-0

40. Tian Y, Cox MA, Kahan SM, Ingram JT, Bakshi RK, Zajac AJ (2016) A Context-Dependent Role for IL-21 in Modulating the Differentiation, Distribution, and Abundance of Effector and Memory CD8 T Cell Subsets. J Immunol. 196: 2153–66. doi: 10.4049/jimmunol.1401236

41. Rosenberg SA, Sherry RM, Morton KE et al. (2005) Tumor progression can occur despite the induction of very high levels of self/tumor antigen-specific CD8+ T cells in patients with melanoma. J Immunol. 175: 6169–76.

42. Gros A, Parkhurst MR, Tran E et al. (2016) Prospective identification of neoantigen-specific lymphocytes in the peripheral blood of melanoma patients. Nat Med. 22: 433–8. doi: 10.1038/nm.4051

43. Restifo NP, Dudley ME, Rosenberg SA (2012) Adoptive immunotherapy for cancer: harnessing the T cell response. Nat Rev Immunol. 12: 269–81. doi: 10.1038/nri3191nri3191[pii]

44. Guo J, Zhang M, Wang B, Yuan Z, Guo Z, Chen T, Yu Y, Qin Z, Cao X (2003) Fractalkine transgene induces T-cell-dependent antitumor immunity through chemoattraction and activation of dendritic cells. Int J Cancer. 103: 212–20. doi: 10.1002/ijc.10816

45. Siddiqui I, Erreni M, van Brakel M, Debets R, Allavena P (2016) Enhanced recruitment of genetically modified CX3CR1-positive human T cells into Fractalkine/CX3CL1 expressing tumors: importance of the chemokine gradient. Journal for immunotherapy of cancer. 4: 21. doi: 10.1186/s40425-016-0125-1

